# Neuroinflammation and functional connectivity in Alzheimer’s disease: interactive influences on cognitive performance

**DOI:** 10.1101/532291

**Authors:** L. Passamonti, K.A. Tsvetanov, P.S. Jones, W.R. Bevan-Jones, R. Arnold, R.J. Borchert, E. Mak, L. Su, J.T. O’Brien, J.B. Rowe

## Abstract

Neuroinflammation is a key part of the etio-pathogenesis of Alzheimer’s disease. We test the relationship between neuroinflammation and the disruption of functional connectivity in large-scale networks, and their joint influence on cognitive impairment.

We combined [^11^C]PK11195 positron emission tomography (PET) and resting-state functional magnetic resonance imaging (rs-fMRI) in 28 humans (13 females/15 males) with clinical diagnosis of probable Alzheimer’s disease or mild cognitive impairment with positive PET biomarker for amyloid, and 14 age-, sex-, and education-matched healthy humans (8 females/6 males). Source-based ‘inflammetry’ was used to extract principal components of [^11^C]PK11195 PET signal variance across all participants. rs-fMRI data were pre-processed via independent component analyses to classify neuronal and non-neuronal signals. Multiple linear regression models identified sources of signal co-variance between neuroinflammation and brain connectivity profiles, in relation to group and cognitive status.

Patients showed significantly higher [^11^C]PK11195 binding relative to controls, in a distributed spatial pattern including the hippocampus, medial, and inferior temporal cortex. Patients with enhanced loading on this [^11^C]PK11195 binding distribution displayed diffuse abnormal functional connectivity. The expression of a stronger association between such abnormal connectivity and higher levels of neuroinflammation correlated with worse cognitive deficits.

Our study suggests that neuroinflammation relates to the pathophysiological changes in network function that underlie cognitive deficits in Alzheimer’s disease. Neuroinflammation, and its association with functionally-relevant reorganisation of brain networks, is proposed as a target for emerging immuno-therapeutic strategies aimed at preventing or slowing the emergence of dementia.

**Significance Statement:** Neuroinflammation is an important aspect of Alzheimer’s disease (AD), but it was not known whether the influence of neuroinflammation on brain network function in humans was important for cognitive deficit.

Our study provides clear evidence that *in vivo* neuroinflammation in AD impairs large-scale network connectivity; and that the link between inflammation and functional network connectivity is relevant to cognitive impairment.

We suggest that future studies should address how neuroinflammation relates to network function as AD progresses; and whether the neuroinflammation in AD is reversible, as the basis of immunotherapeutic strategies to slow the progression of AD.

## Introduction

Neuroinflammation plays a key role in the etio-pathogenesis of Alzheimer’s disease and other neurodegenerative disorders (Edison et al., 2008; Fernandez-Botran et al., 2011; Fan et al., 2015b; Stefaniak and O’Brien, 2016). Pre-clinical models (Heppner et al., 2015; Hoeijmakers et al., 2016; Villegas-Llerena et al., 2016; Li et al., 2018; Wang et al., 2018), and research in humans (Fernandez-Botran et al., 2011; Edison et al., 2013; Fan et al., 2015b; Stefaniak and O’Brien, 2016), demonstrate that microglia of the brain’s innate immune system are activated in Alzheimer’s and related diseases. In addition, genetic association studies have demonstrated a link between Alzheimer’s disease and polymorphisms and mutations of genes linked to immune responses (Villegas-Llerena et al., 2016). Although the mechanisms and mediators of inflammatory risk in Alzheimer’s disease are not fully understood, synaptic and neuronal injury may arise from the release of cytokines and pro-inflammatory molecules such as interleukin-1ß and TGF-ß (Fernandez-Botran et al., 2011), or direct microglial injury to synapses (Hong et al., 2016; Hong and Stevens, 2016). These, in turn, impair synaptic function, network communication, and may accelerate neurodegeneration and synaptic loss (Heppner et al., 2015; Hoeijmakers et al., 2016; Villegas-Llerena et al., 2016; Li et al., 2018; Wang et al., 2018).

Clinical studies of neuroinflammation in dementia have exploited positron emission tomography (PET) ligands that bind to the mitochondrial translocator protein (TSPO) in activated microglia (Cagnin et al., 2001; Gerhard et al., 2006b; Gerhard et al., 2006a; Edison et al., 2008; Edison et al., 2013; Fan et al., 2015b; Fan et al., 2015a; Passamonti et al., In Press). For example, relative to controls, patients with Alzheimer’s disease have higher [^11^C]PK11195 binding in the hippocampus, other medial-temporal lobe regions, and posterior cortices such as the pre-cuneus, which in turn correlates with cognitive severity (Passamonti et al., In Press).

These findings raise the possibility of immunotherapeutic strategies to prevent or slow the progression of Alzheimer’s disease. Nevertheless, key issues remain to be resolved before such therapeutic strategies can be realised. For example, it is necessary to show *how* neuroinflammation is linked to cognitive deficits. A critical and unanswered question is whether regional neuroinflammation changes the functional connectivity of large-scale networks. Such large-scale neural networks represent an intermediate phenotypic expression of pathology in many diseases, that can be non-invasively quantified with resting-state functional magnetic resonance imaging. A challenge is that neither the anatomical substrates of cognition nor the targets of neurodegenerative disease are mediated by single brain regions: they are in contrast distributed across multi-variate and interactive networks.

We thus undertook a multi-modal and multi-variate neuroimaging study to combine [^11^C]PK11195 quantification of distributed neuroinflammation with resting-state functional imaging in patients at different stages of Alzheimer’s disease. We used “source-based inflammetry” (analogous to ‘volumetry’) to reduce the dimensionality (i.e., complexity) of the neuroinflammation signal, and employed multiple linear regression models to associate neuroinflammation, functional network connectivity components, and cognition.

### We tested two key hypotheses

1) that spatially distributed neuroinflammation related to significant changes in large-scale functional connectivity in patients with Alzheimer’s disease, relative to controls.
2) that the relationship between neuroinflammation and abnormal functional connectivity determines cognitive deficit in Alzheimer’s disease.

## Participants & Methods

### Participants

The study was conducted in the context of the Neuroimaging of Inflammation in MemoRy and Other Disorders (NIMROD) study (Bevan-Jones et al., 2017). We included 14 patients meeting clinical diagnostic criteria for probable Alzheimer’s disease (McKhann et al., 2011), and 14 patients with mild cognitive (MCI) patients (15 males and 13 females in total) defined by: i) a mini-mental score examination >24/30; ii) memory impairment at least 1.5 standard deviations below that expected for age and education (Petersen et al., 1999); iii) biomarker evidence of amyloid pathology (positive Pittsburgh Compound-B PET scan) (MCI+) (Okello et al., 2009). We combined patients with clinical Alzheimer’s disease and MCI+ on the basis that these two groups represent a continuum of the same clinical spectrum (Okello et al., 2009).

Fourteen age-, sex-, and education-matched healthy controls (6 males and 8 females) were recruited with no history of major psychiatric or neurological illnesses, head injury or any other significant medical co-morbidity. All participants were aged over 50 years, with premorbid proficiency in English for cognitive testing, did not have any acute infectious or chronic symptomatic systemic inflammatory disorder (e.g., lupus, rheumatoid arthritis, etc.), or contra-indications to magnetic resonance imaging. Patients were identified from the Cambridge University Hospitals NHS Trust Memory Clinics and the Dementias and Neurodegenerative Diseases Research Network (DeNDRoN), while healthy controls were recruited via a Clinical Research Network (http://www.nihr.ac.uk/nihr-in-your-area/dementias-and-neurodegeneration/). Participants had mental capacity and gave written consent in accordance with the Declaration of Helsinki. The study was approved by the local research ethics committee.

### Clinical and cognitive assessment

Clinical indices of cognitive decline included Mini Mental State Examination (MMSE), Addenbrooke’s Cognitive Examination-Revised (ACE-R), and Rey auditory verbal learning test (RAVLT). The demographic and neuropsychological measures are reported in **Table 1**. A Principal Component Analysis on the total MMSE, ACE-R, and RAVLT scores was conducted to reduce the dimensionality of the cognitive deficit into one latent variable which summarized the largest portion of shared variance as the first principal component.

**Table 1.** Participant details (mean, with standard deviation (SD) and range in parentheses) and group differences by chi-squared test, one-way analysis of variance or independent samples t-test. AD/MCI+: Alzheimer’s disease/mild cognitive impairment (amyloid positive on Pittsburgh Compound-B positron emission tomography scan); MMSE: Mini Mental State Examination; ACE-R: Addenbrooke’s Cognitive Examination Revised, RAVLT: Rey Auditory-Verbal Learning Test (delayed recall). NS, not significant with p >0.05 (uncorrected).

| Demographic & Clinical data | AD / MCI+ (N = 28) | Controls (N = 14) | Group difference |
| --- | --- | --- | --- |
| Sex (males/females) | 15/13 | 6/8 | NS |
| Age (years) (SD, range) | 72.3 (±8.6, 53-86) | 68.2 (±5.4, 59-81) | NS |
| Education (years) (SD, range) | 13.0 (±3.0, 10-19) | 14.1 (±2.7, 10-19) | NS |
| MMSE (SD, range) | 25.3 (±2.5, 19-30) | 28.8 (±1.0, 27-30) | T=-4.7, P<0.001 |
| ACE-R (SD, range) | 78.0 (±8.9, 53-91) | 91.5 (±5.3, 79-99) | T=-5.2, P<0.001 |
| RAVLT (SD, range) | 1.4 (±1.6, 0-6) | 9.6 (±3.2, 3-15) | T=-11.0, P<0.0001 |

### Experimental design

#### Structural and functional magnetic resonance imaging protocols and pre-processing

Structural and functional magnetic resonance imaging was performed using a 3 Tesla Siemens Tim Trio scanner with a 32-channel phased-array head coil. A T1-weighted magnetization-prepared rapid gradient-echo image was acquired with repetition time=2300 ms, echo time=2.98 ms, matrix=256×240, in-plane resolution of 1×1 mm, 176 slices of 1 mm thickness, inversion time=900ms and flip angle=9 degrees. The co-registered T1 images were used in a multi-channel segmentation to extract probabilistic maps of 6 tissue classes: grey-matter, white-matter, cerebrospinal fluid, bone, soft tissue, and residual noise. The native-space grey-matter and white-matter images were submitted to diffeomorphic registration to create group template images (Ashburner, 2007). The template was normalised to the Montreal Neurological Institute (MNI) template using a 12-parameter affine transformation. After applying the normalisation parameters from the T1 stream to warp pre-processed functional images into MNI space, the normalised images were smoothed using an 8-mm Gaussian kernel. An estimate of total grey-matter, used in between-subject analysis as a covariate of no interest, was calculated as the median grey-matter tissue intensity in a group mask based on voxels with grey-matter tissue probability of 0.3 across all individuals. Resting state multi-echo functional imaging was carried out for 11 min. A total of 269 echo-planar image volumes were acquired with repetition time=2430ms, echo times=13.00, 30.55 and 48.10 ms, matrix=64×64, in-plane resolution of 3.75×3.75 mm, 34 slices of 3.8 mm thickness with an inter-slice gap of 0.38 mm, and a GeneRalized Autocalibrating Partial Parallel Acquisition (GRAPPA) imaging with an acceleration factor of 2 and bandwidth=2368 Hz/pixel. The first six volumes were discarded to eliminate saturation effects and achieve steady-state magnetization. Pre-processing of resting-state data employed the Multi-Echo Independent Components Analysis (ME-ICA) pipeline, which uses independent component analysis to classify blood oxygenation dependant (BOLD) and non-BOLD signals based on the identification of linearly dependent and independent echo-time related components (https://wiki.cam.ac.uk/bmuwiki/MEICA) (Kundu et al., 2013). This provides an optimal approach to correct for movement-related and non-neuronal signals, and is therefore particularly suited to our study, in which systematic differences in head position might have been expected between groups. After ME-ICA, the data were smoothed with 5.9 mm full-width half maximum kernel.

The location of the key cortical regions in each network was identified by spatial independent component analysis (ICA) using the Group ICA of fMRI Toolbox (Calhoun et al., 2001) in an independent dataset of 298 age-matched healthy individuals from the population-based cohort in the Cambridge Centre for Ageing and Neuroscience (Cam-CAN) (Shafto et al., 2014). Details about pre-processing and node definition are published previously (Tsvetanov et al., 2016). Four networks were identified by spatially matching to pre-existing templates (Shirer et al., 2012). The default mode network contained five nodes: the ventromedial prefrontal cingulate cortex, dorsal and ventral posterior conjugate cortex, and right and left inferior parietal lobes. The fronto-parietal network was defined using bilateral superior frontal gyrus and angular gyrus. Subcortical nodes included nodes having differential group accumulation of [^11^C]PK11195, namely, bilateral putamen and hippocampus. The node time-series were defined as the first principal component resulting from the singular value decomposition of voxels in a 8-mm radius sphere, which was centred on the peak voxel for each node (Tsvetanov et al., 2016).

After extracting nodal time-series we sought to reduce the effects of noise confounds on functional connectivity effects of node time-series using a general linear model (Geerligs et al., 2017). This model included linear trends, expansions of realignment parameters, as well as average signal in the white-matter and cerebrospinal, including their derivative and quadratic regressors from the time-courses of each node (Satterthwaite et al., 2013). The signals in the white-matter and cerebrospinal fluid were created by using the average across all voxels with corresponding tissue probability larger than 0.7 in associated tissue probability maps available in the SPM12 software (http://www.fil.ion.ucl.ac.uk/spm/software/spm12/). A band-pass filter (0.0078-0.1 Hz) was implemented by including a discrete cosine transform set in the general linear model, ensuring that nuisance regression and filtering were performed simultaneously (Hallquist et al., 2013) (Lindquist et al., 2018). The general linear model excluded the initial five volumes to allow for signal equilibration. The total head motion for each participant, which was used in subsequent between-subject analysis as a covariate of no interest (Geerligs et al., 2017), was quantified using the approach reported in Jenkinson and colleagues (Jenkinson et al., 2002), i.e. the root mean square of volume-to-volume displacement. Finally, the functional connectivity between each pair of nodes was computed using Pearson’s correlation on post-processed time-series.

### Positron emission tomography (PET) protocols and pre-processing

All participants underwent [^11^C]PK11195 PET imaging to assess the extent and distribution of neuroinflammation while patients with mild cognitive impairment (MCI) also underwent [^11^C]PiB PET scanning to evaluate the degree of beta-amyloid. [^11^C]PK11195 and [^11^C]PiB PET were produced with high radiochemical purity (>95%), with [^11^C]PiB PET having a specific activity >150 GBq/μmol at the end of synthesis, while [^11^C]PK11195 specific activity was around 85 GBq/μmol at the end of synthesis. PET scanning used a General Electric (GE) Advance PET scanner (GE Healthcare, Waukesha, WI) and a GE Discovery 690 PET/CT, with attenuation correction provided by a 15min 68Ge/68Ga transmission scan and a low dose computed tomography scan, respectively. The emission protocols were 550 MBq [11C]PiB injection followed by imaging from 40-70 minutes post-injection, and 75 minutes of dynamic imaging (55 frames) starting concurrently with a 500 MBq [^11^C]PK11195 injection. Each emission frame was reconstructed using the PROMIS 3-dimensional filtered back projection algorithm into a 128×128 matrix 30cm trans-axial field of view, with a trans-axial Hann filter cut-off at the Nyquist frequency (Kinahan and Rogers, 1989). Corrections were applied for randoms, dead time, normalization, scatter, attenuation, and sensitivity.

For [^11^C]PiB we used reference tissue region of interest (ROI) defined by ≥ 90% on the SPM8 grey-matter probability map (smoothed to PET resolution) in the cerebellar cortex (Schuitemaker et al., 2007). For [^11^C]PK11195, supervised cluster analysis was used to determine the reference tissue time-activity curve (Turkheimer et al., 2007). [^11^C]PiB data were quantified using standardized uptake value ratio (SUVR) by dividing the mean cerebrospinal fluid (CSF) corrected radioactivity concentration in each Hammers atlas ROI by the corresponding mean CSF-corrected radioactivity concentration in the reference tissue ROI (whole cerebellum). [^11^C]PiB data were treated as dichotomous measures (i.e., positive or negative) and considered positive if the average SUVR value across the cortical ROIs was > 1.5 (Hatashita and Yamasaki, 2010). For [^11^C]PK11195 maps of non-displaceable binding potential (BP_ND_), a measure of specific binding, were determined using a basis function implementation of the simplified reference tissue model, both with and without CSF contamination correction (Gunn et al., 1997). [^11^C]PK11195 BP_ND_ maps (termed here PK maps for simplicity) were also generated using this basis function approach.

The PK maps were co-registered and warped to the Montreal Neurological Institute (MNI) space using the flow fields. To minimise the noise effects from non grey-matter regions, the normalised PK maps were masked with a group-based grey-matter mask based on voxels having grey-matter tissue probability larger than 0.3 in grey-matter segmented images across all individuals. The normalised images were smoothed using a 6mm Gaussian kernel. We then used independent component analysis across participants to derive spatial patterns of PK maps across voxels expressed by the group in a small number of independent components. All PK maps were spatially concatenated and submitted to Source-Based ‘inflammetry’ (SBI) to decompose images across all individuals in a set of spatially independent sources without providing any information about the group (Xu et al., 2009), using the GIFT toolbox. Specifically, the *n*-by-*m* matrix of participants-by-voxels was decomposed into: (i) a source matrix that maps each independent component to voxels (here referred to as PK_IC_ maps), and (ii) a mixing matrix that maps PK_ICs_ to participants. The mixing matrix consists of loading values (one per participant) indicating the degree to which a participant expresses a defined PK_IC_. The independent component loading values for the PK_IC_ were taken forward to between-participant analysis of functional connectivity (**Figure 1**), if they were (a) differentially expressed by controls and patients with Alzheimer’s disease pathology; and (b) were associated with atrophy (see results section and Fig.3). Only 1 dependent variable (IC3) met these criteria.

**Figure 1.**
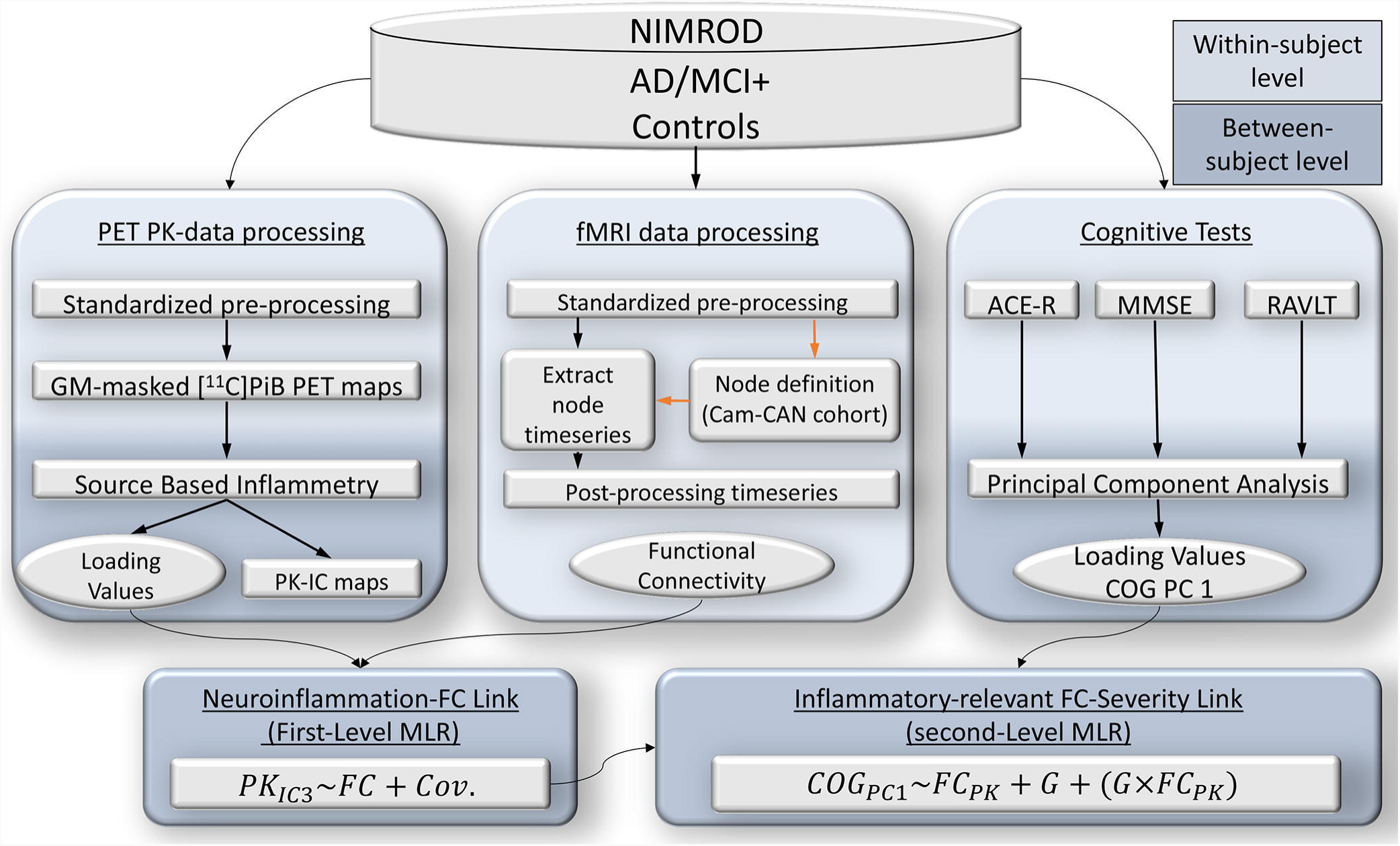
Schematic representation of various modality datasets in the study, their processing pipelines on a within-subject level (light blue), as well as data-reduction techniques and statistical strategy on between-subject level (dark blue) to test for associations between the datasets. Abbreviations: PK_IC_ (Independent component [^11^C]PK11195 maps); FC (functional connectivity); Cov. (covariates); COG PC1 (latent variable (cognitive deficit) which summarizes the largest portion of shared variance as the first principal component); MMSE (Mini Mental State Examination), ACE-R (Addenbrooke’s Cognitive Examination-Revised); RAVLT (Rey auditory verbal learning test); GM (grey-matter); PiB (Pittsburgh Compound-B positron emission tomography); Cam-CAN (Cambridge Centre for Ageing and Neuroscience); AD/MCI+ (Alzheimer’s disease and MCI PiB positive mild cognitive impairment patients); MLR (multiple linear regression analyses); NIMROD (Neuroimaging of Inflammation in MemoRy and Other Disorders study).

### Statistical analyses

We adopted a two-level procedure, in which, at the first-level, we sought to identify functional connectivity differences associated with differences in [^11^C]PK11195 binding. In a second-level analysis, we tested whether individual variability in functional connectivity (from first-level analysis) is specifically associated with variability in cognitive decline in the group of patients with Alzheimer’s disease pathology.

Details about the first-level analysis approach are published previously (Tsvetanov et al., 2018). In short, we used multiple linear regression with well-conditioned shrinkage regularization (Ledoit and Wolf, 2004) to identify correlated structured sources of variance between functional connectivity and neuroinflammation measures. In particular, this analysis describes the linear relationship between functional connectivity and PK maps on a between-subject level, in terms of structure coefficients (Thompson and Borrello, 1985), by providing a linear combination of the functional connectivity measures, which we term *brain scores*, that are optimised to be highly correlated with the between-subject variability in the expression of the PK maps. Namely, brain-wide connectivity strength for each individual defined the independent variables, and PK_IC_ subject-specific loading values for group differentiating components were employed as dependent variable.

To identify and exclude potential outliers, Grubbs’ test was used (Grubbs, 1969; Barnett and Lewis, 1994). None of the loading values in the IC3 were outlying observations. Furthermore, to down-weight the effects of extreme or imprecise data points, analyses used robust linear regression. To avoid overfitting, first-level multiple linear regression model was integrated with a 5-Fold Cross–Validation (Thompson and Borrello, 1985). To minimize the non-negligible variance of traditional k-Fold cross-validation procedure, we repeated each k-Fold 1,000 times with random partitioning of the folds to produce an R-value distribution, for which we report the median values.

Next, we tested the hypothesis that the effect of neuroinflammation on functional connectivity was related to cognitive deficits, in patients relative to controls. To this end, we performed a second-level multiple liner regression (MLR) analysis. Independent variables included subjects’ *brain scores* from first level MLR (reflecting how strongly each individual expressed the whole brain pattern of functional connections weighted by the IC3-PET derived data), group information, and their interaction term (brain scores x group). The dependent variable was subjects’ loading values of the first principal component across the three cognitive tests. Covariates of no interest included age, gender, head movement, and global grey-matter volume.

## Results

### Source-based ‘inflammetry’

The optimal number of components (n=5) was detected with minimum-distance length criteria. One component showed significant differences between the patient and control groups in terms of their loading values (PK_IC3_, t-value = −2.1, p-value = 0.04) (Figure 2 right panel). The spatial extent of this PK_IC3_ included voxels with high values in cortical and sub-cortical regions, including the inferior temporal cortex and hippocampus, indicating that individuals with higher loading values, in this case the patient group, had higher [^11^C]PK11195 binding in these regions, relative to the control group (Figure 2, left panel). The other components did not differentiate patients from controls (Figure 3, first row). The PK_IC3_ component, which differed between patients and controls, was the only PK component that negatively correlated with total grey-matter values in patients but not controls (Figure 3, second and third row). In other words, the patients expressing higher [^11^C]PK11195 binding showed also higher levels of cortical atrophy (Figure 3, second and third row). This result was obtained when including the grey-matter volume as a covariate of no interest in the analysis, which suggests that the reported association was over and above the effects of overall brain atrophy.

**Figure 2.**
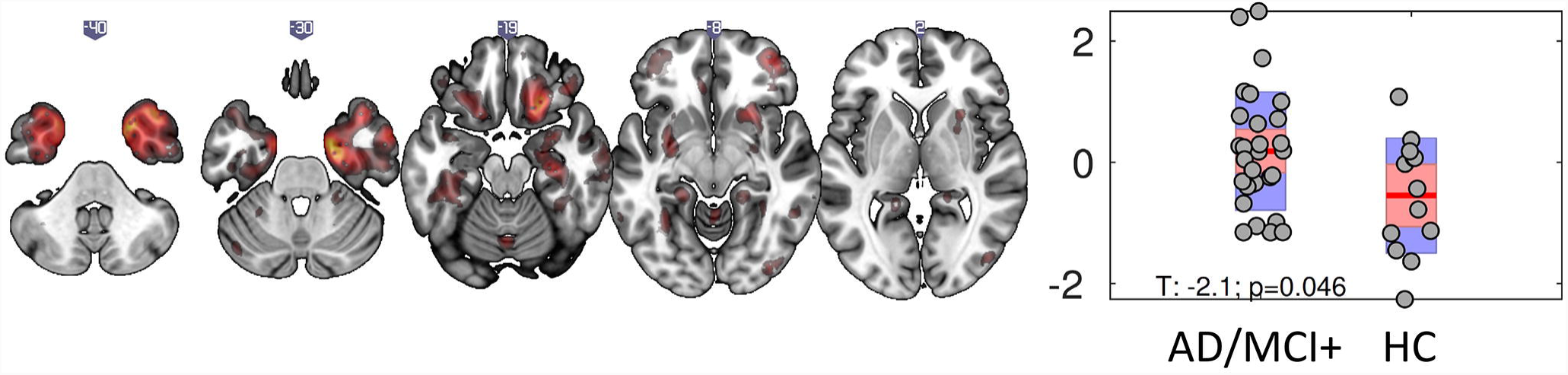
Source-Based Inflammetry for the component differentially expressed between groups: (left) independent component (IC) spatial map reflecting increase in [^11^C]PK11195 (PK) binding values in cortical and subcortical areas including inferior temporal cortex and hippocampus, regionally specific increase over and above global PK differences between groups (regions in red), (right) bar plot of subject loading values for AD/MCI+ and control group (each circle represents an individual) indicating higher loading values for AD/MCI+ than control group as informed by two-sample unpaired permutation test. AD/MCI+ (Alzheimer’s disease and MCI PiB positive mild cognitive impairment patients).

**Figure 3.**
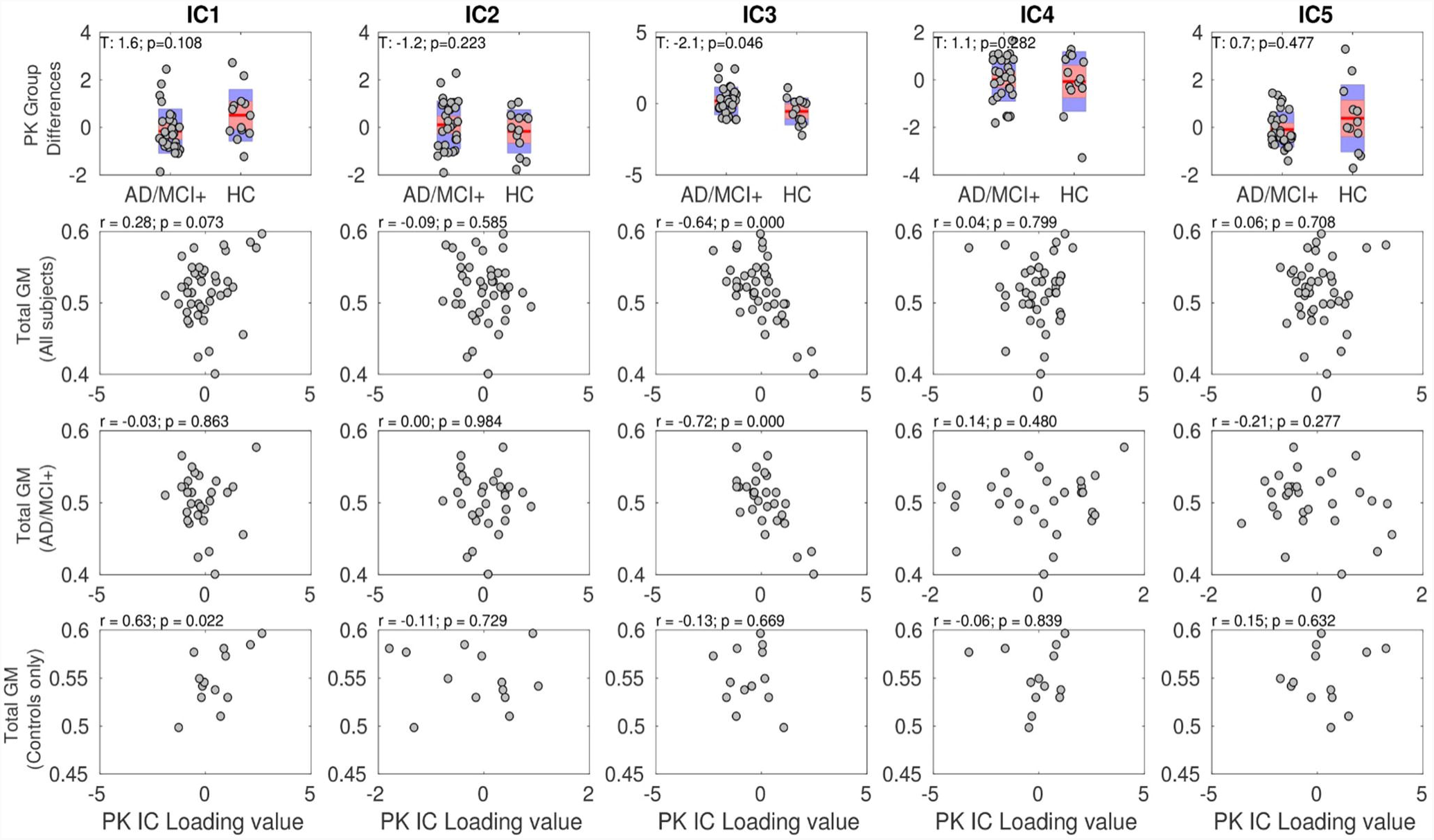
The Source-Based Inflammetry (SBI) identified five independent components (IC) which reflected [^11^C]PK11195 (PK) binding values in cortical and subcortical areas. The PKIC3 component differed between AD/MCI+ patients and controls (first row, third column). This PKIC3 component negatively correlated with total grey-matter volumes in all individuals as well as in patients-only (but not controls-only) (second, third, and fourth rows). In other words, the patients expressing higher [^11^C]PK11195 binding PKIC3 component (reflecting higher binding in the inferior temporal cortex and hippocampus as shown in Figure 2) also displayed higher levels of brain-wide atrophy.

All in all, our findings imply that the PK_IC3_ component reflects specific patterns of neuroinflammation and neurodegeneration in Alzheimer’s disease. These patterns were next tested in terms of their relevance for changes in large-scale network function and their interactive effect in predicting cognitive deficit in Alzheimer’s disease.

### Functional connectivity

As expected, there was strong positive functional connectivity between all nodes within the four networks, identified by spatially matching to pre-existing templates (Figure 4, left panel). In terms of group differences, the functional connectivity within networks (within the default mode network, within the fronto-parietal network, left-right putamen and left-right hippocampus) and between the default mode network and hippocampus was weaker in patients relative to controls (Figure 4, right panel). Furthermore, the connectivity between the putamen and hippocampus increased while the connectivity between default mode network and putamen was less negative for patients relative to controls.

**Figure 4.**
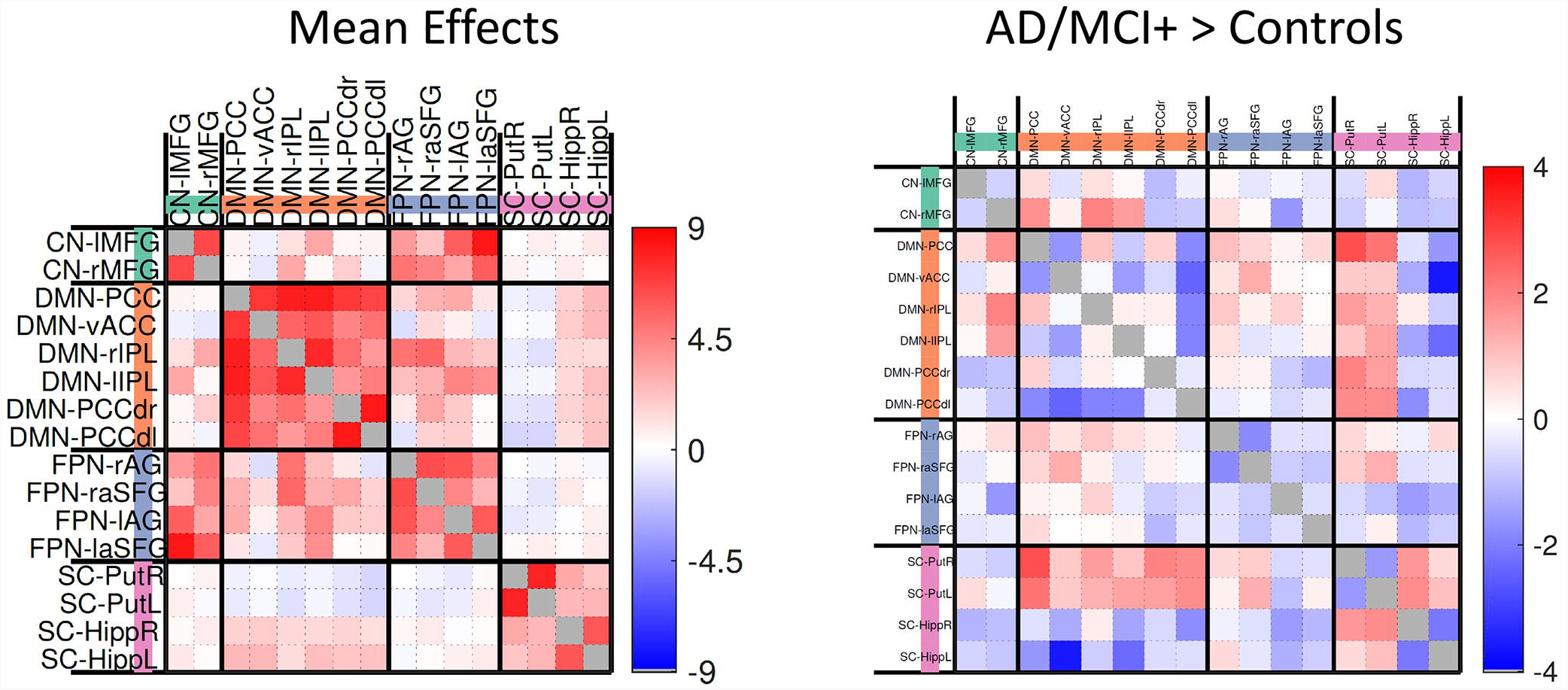
Main effects (left) and group difference effects (AD/MCI+>Controls, right) between default mode network (DMN) and subcortical regions using univariate approach. ACC – anterior cingulate cortex; PCC – posterior cingulate cortex; IPL – intraparietal lobule; FPN – fronto-parietal network; Put – Putamen; Hipp – Hippocampus, AG – angular gyrus; SFG – superior frontal gyrus; R, right; L, left. Note that the whole pattern of brain connectivity rather than each connection separately was employed to study how subject-specific neuroinflammatory levels influence large-scale network connectivity (Fig 5).

### Functional connectivity and neuroinflammation

The first-level multiple linear regression model assessing the relationship between PK_IC3_ maps and functional connectivity data was significant (r=0.52, p<0.001). The standard coefficients indicated a positive association between the PK_IC3_ loading values and variability in functional connectivity (Figure 5, left panel). In other words, individuals with higher [^11^C]PK11195 binding values in the inferior temporal cortex and medial temporal lobe regions (as reflected by higher PK_IC3_ values) showed: i) increased connectivity between the default mode network, the hippocampus, and other subcortical regions, and ii) weaker connectivity for nodes within the default mode network.

**Figure 5.**
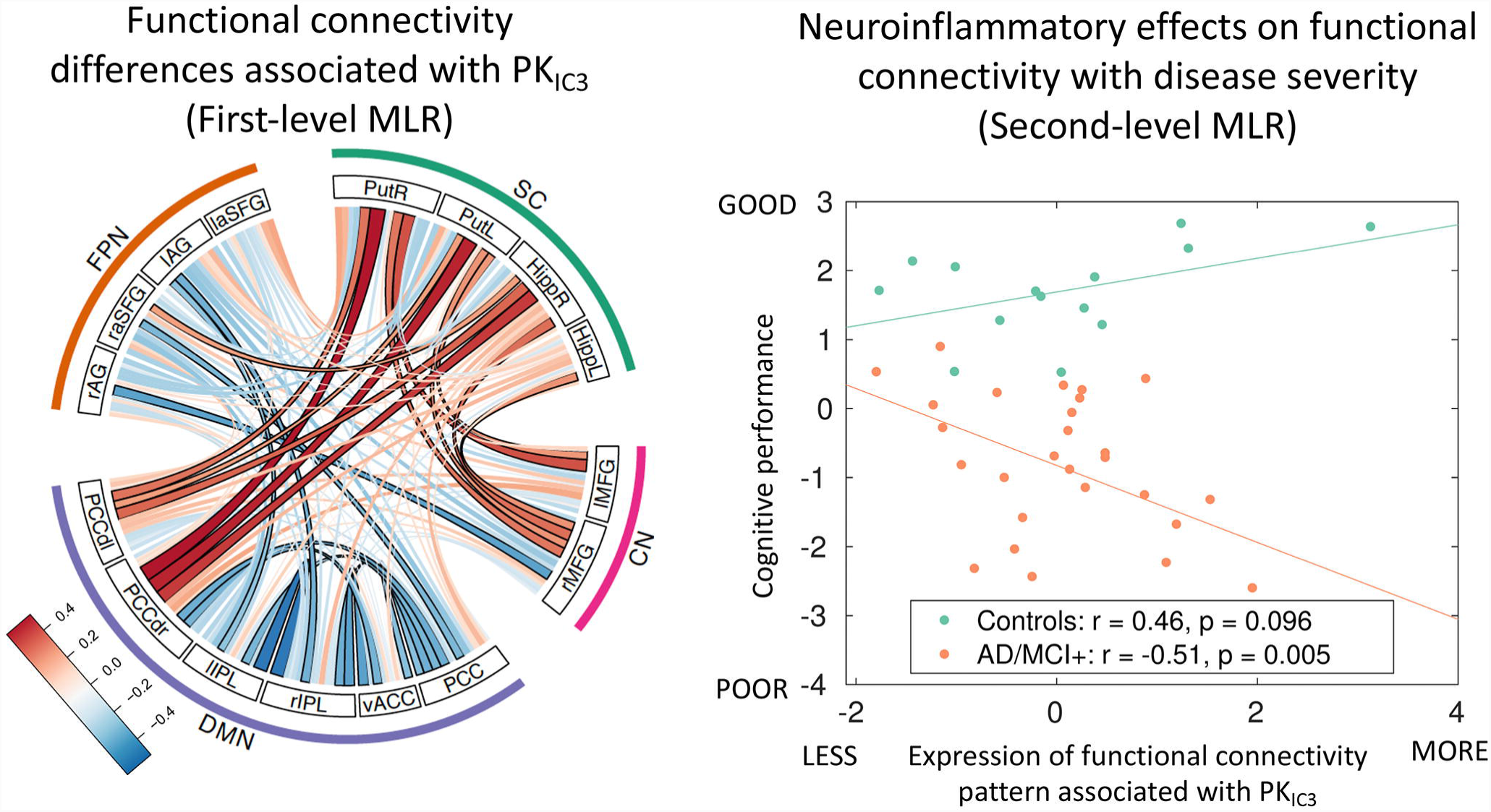
(Left) First-level multiple linear regression (MLR) indicating that functional connectivity differences (deviating from groups effects in Figure 3) are associated positively with [^11^C]PK11195-related independent component measures (PK_IC3_). Connections surviving a threshold of P<0.05 corrected for multiple comparisons are highlighted with a black contour, although it is important to bear in mind that the whole-pattern of brain connectivity was used in the analysis shown in the right panel. (Right) Second-level MLR association between PK_IC3_ pattern of functional connectivity and cognitive performance for patients with Alzheimer’s disease (AD) pathology (including mild cognitive impairments, amyloid positive, MCI+) (orange) and control (green) groups. The group difference in slopes was significant (p<0.0001). ACC – anterior cingulate cortex; PCC – posterior cingulate cortex; IPL – intraparietal lobule; FPN – fronto-parietal network; SC – subcortical, DMN – default mode network, DMNd – dorsal DMN, Put – Putamen; Hipp – Hippocampus, AG – angular gyrus; SFG – superior frontal gyrus; R, right; L, left.

### Linking neuroinflammation, connectivity, and cognitive deficits

The first component of the principal component analysis (PCA) of cognitive tests explained the 80% of the variance across the three cognitive measures (with coefficients .61, .61, and .52 for MMSE, ACE-R, and RAVLT, respectively).

Next, we tested whether the effects of neuroinflammation on network connectivity were specific to the patient group and whether this functionally-relevant neuroinflammation related to cognitive deficits. Consistently with this hypothesis, the interaction term between group and *brain scores* (reflecting how strongly each individual expresses the pattern shown in Fig.5 (which is the brain-wide pattern of functional connections optimised to highly correlate with the IC3-PET derived data) was significantly associated to the first component of the PCA of cognitive tests (t=-3.4, p=0.004).

A *post-hoc* analysis within each group indicated a significant negative association between the behavioural scores from the PCA and functional connectivity/PK-combined indices in the patient group (r= −0.51, p=0.005) (Figure 5, right panel). Conversely, a non-significant positive direction of association for the same relationship between PCA-derived cognitive scores and brain measures was found in controls (r=0.46, p=0.09). The significant difference between patients and controls remains if AD and MCI+ subgroups are analysed separately (not shown). The negative association in the patient group indicated that patients in whom higher neuroinflammation was more strongly associated with more abnormal connectivity also performed worse on a summary measure of cognitive deficits.

## Discussion

This study establishes a link between the presence of neuroinflammation and the disruption of large-scale functional connectivity in Alzheimer’s disease. The degree to which patients expressed the association between abnormal connectivity and neuroinflammation *itself* correlated with their cognitive deficit. This relationship was found across the spectrum of patients ranging from mild cognitive impairment with biomarker evidence of Alzheimer’s pathology to clinical diagnosis of probable Alzheimer’s disease. We suggest that not only does neuroinflammation relate to large-scale network function, but that the disruption of connectivity linked to neuroinflammation mediates cognitive deficits in Alzheimer’s disease.

There are different mechanisms by which neuroinflammation might alter brain functional connectivity and consequently neuroimaging indices of network function or *vice versa* (i.e., the ways in which synaptic firing can influence microglia activity and related neuroinflammatory processes). Microglia are important contributors in the process of synaptic pruning and regulation of synaptic function (Hong and Stevens, 2016). The microglia’s highly mobile and ramified branches can reach and surround synaptic terminals to promote phagocytosis and synaptic demise (Hong and Stevens, 2016). Microglia-induced complement activation might also contribute to synaptic dysfunction and loss, especially in the context of amyloid deposition and neuritic plaque formation (Hong et al., 2016). On the other hand, synaptic firing can influence microglia activation via specific membrane receptors and ion channels (Tofaris and Buckley, 2018).

The anatomical distribution of neuroinflammation in Alzheimer’s disease and its effects on large-scale network function supports the hypothesis that neuroinflammation might be an early event in the pathogenesis of Alzheimer’s disease and that our current results are not driven by a global effect of a systemic inflammatory confound which would have affected the whole-brain indistinctively.

This study has also two important implications. First, it reinforces the notion that neuroinflammation is a key pathophysiological mediator of Alzheimer’s disease and its clinical variability (Weiler et al., 2016). Genome-wide association studies have challenged the idea that neuroinflammation is merely a secondary event caused by neurodegeneration and have conversely sustained a primary role of microglia-related molecular pathways in the etio-pathogenesis of Alzheimer’s disease (Guerreiro et al., 2013; Jonsson et al., 2013). For instance, mutations in TREM2, an immune cells receptor expressed on microglia, represent a risk factor for Alzheimer’s disease and other neurodegenerative disorders (Guerreiro et al., 2013; Jonsson et al., 2013). Together with our results, these data suggest that immunotherapeutic strategies might be helpful to reduce the deleterious impact of neuroinflammation on cognitive deficit in Alzheimer’s disease.

Second, the functional connectivity abnormalities observed here can be considered an intermediate phenotypic expression of the neuroinflammatory pathology in Alzheimer’s disease. This can be relevant to reconcile the apparent conflict between the encouraging findings from basic research on the role of neuroinflammation in Alzheimer’s pathogenesis (Heppner et al., 2015) and the results from human studies which as yet have provided little support for immunotherapeutics in Alzheimer’s disease (Group et al., 2007; Group et al., 2008), despite epidemiological evidence (Breitner and Zandi, 2001; in t’ Veld et al., 2001).

In other words, assessing how neuroinflammation influences the intermediate phenotypes of large-scale network functional connectivity might help explaining why clinical trials have failed thus far to demonstrate a role for immunotherapeutic strategies due to high patient heterogeneity. Our data showed marked individual differences in the relationship between resting-state functional connectivity and neuroinflammation in patients with Alzheimer’s disease at different stages, and it was this variance that was significantly related to individual differences in cognitive performance.

Our study has limitations and caveats. First, we recognise that even the multi-variate methods of statistical associations used here do not *in themselves* demonstrate causality between neuroinflammation, network dysfunction, and cognition. To address this issue, longitudinal and interventional studies are needed, alongside mediation analyses (Fan et al., 2015a; Kreisl et al., 2016).

Second, the molecular pathology of Alzheimer’s disease is multi-faceted, with amyloid deposition, tau accumulation, and vasculopathy. These processes, alone or in combination, may moderate the association between neuroinflammation and functional connectivity; hence, multi-modal studies that capture each of these aspects will be useful to formally assess the complex interplay between neuroinflammation, abnormal tau deposition, vasculopathy, and cognitive deficits.

Third, the confounding effect of head motion on functional imaging has been fully recognized as both challenging and critical for interpretation of functional imaging studies, especially in clinical populations. To minimize this confound, we used Multi-Echo Independent Components Analysis (ME-ICA) and validated pre-processing pipelines, which separate changes in the fMRI signal that are due to blood oxygenation dependant (BOLD) and non-BOLD signals. Furthermore, we included movement-related parameters as covariates of no interest in *second-level* analyses, as well as motion and physiological signals in *first-level* analyses.

Fourth, the use of the [^11^C]PK11195 tracer has its own limitations in terms of reduced affinity to the mitochondrial translocator protein (TSPO) in activated microglia, especially when compared to second-generation TSPO tracers as PBR28 (Fujita et al., 2017). On the other hand, such second-generation TSPO tracers are affected by common genetic polymorphisms (Owen et al., 2012).

Finally, at the phenotypic level, it remains to be determined whether the deleterious impact of neuroinflammation on network function can be revealed in pre-symptomatic adults at risk of Alzheimer’s disease, for example in carriers of autosomal dominant genetic mutations. Despite the inclusion of patients with mild cognitive impairment with biomarker evidence of Alzheimer’s pathology, our study cannot resolve the timing of neuroinflammation and its causal relationship to network dysfunction, cell loss, and cognitive deficit.

In conclusion, we have shown that source-based ‘inflammetry’ of [^11^C]PK11195 PET data reveals a distributed profile of neuroinflammation in Alzheimer’s disease, which in turn related to abnormal functional connectivity. Our cross-modal multivariate analyses also indicated that heterogeneity in cognitive status was associated to variability in neuroinflammation-related network dysfunction. These data emphasize the value of multi-modal neuroimaging to study how different aspects of the molecular pathology of Alzheimer’s disease mediate brain function and cognition. Improved stratification procedures may facilitate more efficient therapeutic trials in Alzheimer’s disease, based not on inflammation, tau, atrophy or connectivity alone, but on their complex interaction that leads to individual differences in cognitive impairment.

## Acknowledgments & Conflict of Interest

We thank our volunteers and the radiographers/technologists at WBIC and PET/CT, Addenbrooke’s Hospital, for their invaluable support in data acquisition. We thank the NIHR Eastern Dementias and Neurodegenerative Diseases Research Network for help with subject recruitment. We thank Dr Istvan Boros and others at WBIC RPU for the manufacture of [^11^C]PK11195 and [^11^C]PiB. This study was funded by the National Institute for Health Research Cambridge Biomedical Research Centre and Biomedical Research Unit in Dementia (NIHR, RG64473), the Wellcome Trust (JBR 103838), and the Medical Research Council (RG91365/SUAG/004 and MR/P01271X/1). Dr. Li Su is supported by Alzheimer’s Research UK. L. Passamonti, K. Tsvetanov, W.R. Bevan-Jones, P.S. Jones, R. Arnold, R. Borchert, E. Mak, and L. Su report no financial disclosures or conflict of interest relevant to the manuscript. J.T. O’Brien has served as deputy editor of International Psychogeriatrics, received grant support from Avid (Lilly), and served as a consultant for Avid and GE Healthcare, all for matters not related to the current study. J.B. Rowe serves as editor to Brain, has been a consultant for Asceneuron and Syncona, and has received academic grant funding from AZ-MedImmune, Janssen, and Lilly, unrelated to this study.

